# The neural analysis toolkit unifies semi-analytical techniques to simplify, understand, and simulate dendrites

**DOI:** 10.1101/2025.06.26.661734

**Authors:** Willem A.M. Wybo

## Abstract

While simulating compartmental dynamics in response to various input patterns is the prevalent technique for understanding dendritic computation, a great deal can be learned from classical analytical methods that provide solutions for the dendritic voltage. For example, such solutions are needed to simplify spatially extended neuron models, to understand frequency-dependent response properties, to elucidate the interaction between synaptic inputs, and hence to reveal the effective compartmentalization of dendrites into functional subunits. Nevertheless, these methods have not been implemented in modern software tools. This works describes the NEural Analysis Toolkit (NEAT), a Python toolbox that implements classical algorithms to compute response properties of spatially extended neuron models, and that leverages these algorithms to simplify them. Packaged with this are a range of useful utilities to plot morphologies and spatial quantities defined on the morphology, to distribute locations on the morphology, and to select parts of the morphology to e.g. apply morphological ablations or alter the membrane properties. The resulting models can be exported to NEURON and NEST, two commonly used simulators, the former focused on detailed single neuron model simulations, and the latter geared towards distributed network simulations. As a consequence, this toolbox provides a missing link between single neuron computation and large-scale network analysis, substantially facilitating the study of the role of dendritic computation in shaping emergent network dynamics. NEAT is available through `pip` under the name `‘nest-neat’`, or from its source code (https://github.com/nest/NEAT), which is provided under the GNU General Public License. Furthermore, support for NEAT is provided on its GitHub page and through the NEST user mailing list.

## 1 Introduction

Computational neuroscience is driven by software tools. To simulate networks of neurons, and understand the effects of synaptic plasticity, computational neuroscientists typically rely on NEURON to investigate morphologically detailed neurons [1], on NEST when the efficiency of network simulations is of essence [2], and on Brian when model customization and ease of implementation is of primary importance [3]. Over the years, a rich ecosystem of models has been developed for each of these tools. In NEURON, the primary use case is spatially extended neuron models, where the membrane is equipped with currents whose conductance depends on the local membrane potential and/or presynaptic transmitter release, and whose dynamics can be made arbitrarily complex through the .mod model description language (see e.g. ModelDB [4]). In NEST, the collection of available neuron models has so far been more simple, with a range of predefined neuron models provided by default, and the recent development of a model-description language which, in its implicit assumptions, was geared towards point neurons [5]. However, the algorithms that transmit spikes between neurons have been honed to enable a highly efficient communication on massively parallel computing systems [6, 7]. Finally, Brian offers much freedom in the model definition, converting arbitrary differential equations in C++ code, but does not offer the advanced parallellisation functionalities to simulate large networks [3]. These simulation tools, therefore, encourage development of specific branches of computational neuroscience, by each creating a niche of applications in which model development is easy, while research that falls outside of these areas of application is typically hard, and requires development of extensive problem-specific research codes.

One of the areas of research that falls outside of the realm of applicability of standard software tools, but that is fundamental towards understanding brain function, is that of understanding the effective degree of dendritic complexity at which brain circuits operate [8–10]. While simulators like NEURON [1, 11] and Arbor [12, 13] implement the discretisation of neural morphologies into coupled ordinary differential equations through the 2nd order finite difference approximation, internally effectively constructing a representation of the neuron model as a series of coupled electrical compartments, they offer no native tools to substantially simplify these models. While many simplified, rate-based neuron models have been proposed that convert the dendrite into multi-layer perceptron-like models [14–17] – leading authors to explore such neuron models in a machine learning context [18–20] – this work focuses on descriptions in terms of electrical compartments. Such description result in spiking models, and offer a biophysical interpretation for their parameters. In a top-down approach, reduced compartmental models are often constructed as combinations of few coupled compartments, whose conductance parameters are optimized to reproduce chosen dendritic computations [21–26]. On the other end of the spectrum, bottom-up approaches start from highly detailed models, simplifying them by requiring that the reduced model conserves geometrical quantities like surface area [27, 28], or electrical properties like attenuation [29] or transfer impedance [30]. Such approaches can essentially be thought of as finding appropriate rescaled conductances for the compartmental currents. Prior work has shown that the inverse of the dendritic resistance matrix, evaluated at dendritic sites of interest, yields precisely the desired conductance parameters [31]. The resulting simplification methodology is useful for both the top-down approach [25] as well as the bottom-up approach [31].

The simplification algorithm by Wybo et al. [31] requires evaluating large numbers of such resistance matrices around different expansion points, to generalize well to non-linear dynamical regimes. Such analytically computable quantities furthermore yield insight into response properties of the neuron. Input and transfer resistance matrices yield insight into the compartmentalisation of the neuron model into functional subunits [32, 33], and in spatial summation across the dendritic tree [34]. Input and transfer impedances yield insight in the neuron’s spatio-temporal responses to different input frequencies [35]. Computing the time-scales of the neural response kernels provides information on the effect of dendritic morphology on input integration and action potential firing [36], while computing conductance loads explains spike feature variability [37] and excitation-inhibition interactions [38]. Standard software tools, however, do not provide analytical algorithms to compute such response properties. While these quantities can be derived approximately from simulation results, they can be computed with more efficiency and/or accuracy through analytical algorithms, such as Koch’s [39] or Abbotts [40–42] algorithm to compute input and transfer resistances and impedances [39], or Major’s algorithm to compute the separation of variables expansion [43–46], which provides response timescales and their associated spatial profiles [47].

Supported by the NEST initiative, this paper presents the NEural Analysis Toolkit (NEST-NEAT), which as it primary application, aims to make neuron simplification easy. In doing so, NEST-NEAT thus provides a bridge between single- and multicompartment modeling. In general, there is a need to make interacting with complex dendrite models easier [48]. NEST-NEAT relies largely on Python, and provides a Pythonic and convenient of way of defining morphologically extended neuron models, reading the commonplace ‘.swc’ format for specifying morphologies into an internal representation of cylindrical sections. NEST-NEAT also provides a number of general routines interacting with the morphology, distributing and defining sets of locations, and for specifying biophysical mechanisms. Furthermore, NEST-NEAT implements Koch’s algorithms to compute input and transfer impedances in the Fourier domain [49], and appropriate custom inverse transform algorithms to convert these frequency-domain impedances to response kernels in the time domain. NEST-NEAT also implements Major’s separation of variables algorithm [43–46], to compute the response timescales of neuron models and their associated spatial profiles. NEST-NEAT uses these analytical algorithms to implement Wybo’s simplification algorithm [31], which can be used to derive accurate simplifications of arbitrary complexity. Finally, NEST-NEAT can be used to export full and reduced models to both NEURON and NEST for simulation.

## 2 Design and implementation

### 2.1 An overview of NEAT

As dendrites are trees, NEST-NEAT (referred to simply as NEAT in the remainder of this text for brevity) is built almost entirely on tree-graph data structures (Fig 1A). A tree-graph is a graph without loops. As such, one node can be understood to be the root of the tree. Each node then stores a number of child nodes, and a reference to its parent (`None` in case of the root). NEAT features three types of tree classes: abstract, compartmental and morphological. The abstract tree class `STree` is the base class from which all trees inherit, and implements basic functionality to iterate through trees, to add and remove nodes, and queries to return specific collections of nodes (e.g. the subtree of a specific node). In the compartment tree classes (`CompartmentTree` and derived classes), each node is assumed to be a single electrical compartment coupled to its parent through a coupling conductance (Fig 1B). Finally, in the morphological tree classes (`MorphTree` and derived classes) each node is assumed to signify a cylindrical section of neurite, which connects the `xyz`-coordinate of the parent node to its own `xyz`-coordinate.

**Figure 1.**
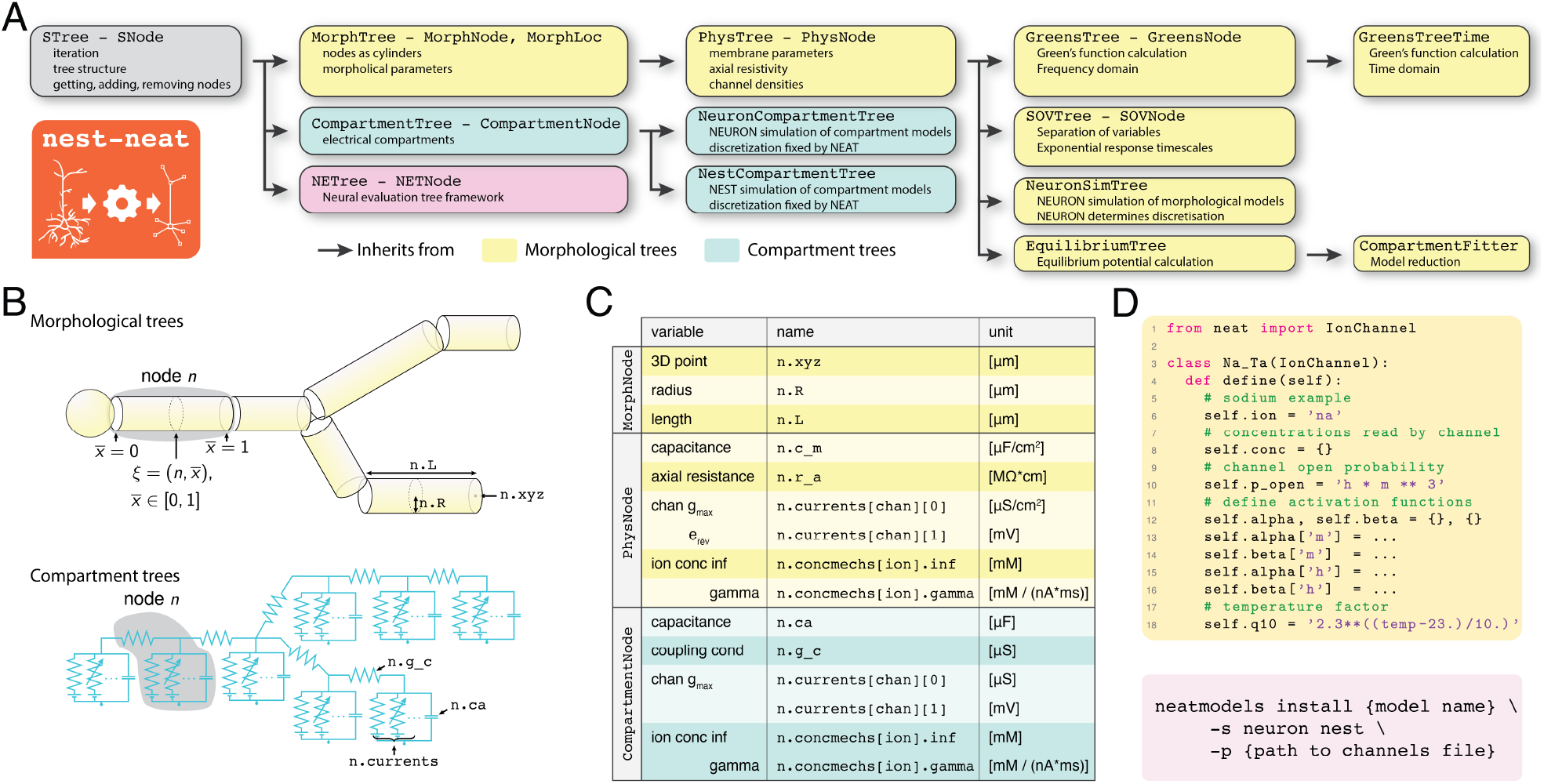
The structure of NEAT. **A:** Overview of the inheritance structure of NEAT’s trees and associated node classes. In morphological trees (yellow), nodes represent cylindrical sections, whereas in compartment trees (blue), nodes represent electrical compartments. The neural evaluation tree (NETree) is a specialty tree developed to analyze independence of subunits [33]. **B:** In morphological trees (top), the node index `n` refers to a cylindrical section with a length `n.L`, radius `n.R` and the xyz-coordinate of the endpoint `n.xyz`. A further coordinate 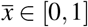 is needed to fully specify a location. In compartment trees (bottom), only a node index `n` is required. A compartment node `n` is coupled to its parent through the coupling conductance `n.g_c`. **C:** Table of critical parameters stored at each node, how to access them in the data structures, and their units. Note that nodes have the same inheritance structure as trees, so that a `PhysNode` inherits from `MorphNode`, and therefore also contains all its parameters. **D:** Code example of an `IonChannel` in NEAT. At present, NEAT only supports Hodgkin-Huxley type channels in its analytical calculations, from which ion channel simulation code for NEURON and NEST can be generated. **E:** To generate simulation code for NEURON and/or NEST, NEAT provides the neatmodels terminal script, which compiles groups of ion channels either into `*.mod`-files or into `*.nestml`-files.

In NEAT, the various tree classes either add a layer of complexity, or implement a different functionality. For instance, NEAT’s `MorphTree` – initialized from the de-facto standard .swc-format [52]– stores the morphology of the neuron, and provides functionality to interact with it, such as by distributing locations on the morphology. However, it does not store physiological parameters; this is implemented by NEAT’s `PhysTree`. The `PhysTree` inherits from `MorphTree`, and adds functionality to set physiological parameters, such as passive parameters (axial resistance, membrane capacitance, leak conductance and reversal) or ion channel densities and concentration mechanics. A `PhysTree` thus represents a detailed biophysical neuron model. A number of classes then inherit from `PhysTree` to implement specific computations, such as the `SOVTree`, which computes the separation of variables expansion through Major’s algorithm, the `GreensTree`, which computes the Green’s function in the Frequency domain through Koch’s algorithm, the `GreensTreeTime`, which transforms frequency domain results to the time domain, the `EquilibriumTree`, which computes the equilibrium potential throughout the tree, and finally the `CompartmentFitter`, which derives simplified neuron models and returns them as `CompartmentTree` instances. As all these trees inherit from `PhysTree`, they can be instantiated using the same routines to specify morphology and physiology. However, if a tree is first built as one specific tree class, it would be cumbersome to construct it in the same way if another computation is needed. To that purpose, NEAT implements a copy-construct functionality, so that any tree type can also be instantiated from any other tree class instance.

Finally, to simulate neuron models, NEAT implements the `NeuronSimTree`, which inherits from `PhysTree` and exports full morphological models (where nodes are combinations of cylinders) to NEURON. NEAT also defines the `NeuronCompartmentTree` and `NestCompartmentTree`, which inherit from `CompartmentTree` and export reduced neuron models to NEURON resp. NEST.

Morphological reconstructions, specified in the .swc format, often feature unbranched sections of dendrite, consisting of many nodes with identical radii, while the physiological parameters of the model are also identical for these nodes. In principle, such a section could be replaced by a single cylinder, without changing the underlying mathematical description. However, NEAT’s `MorphTree` aims to preserve a tree structure that is a one-to-one mapping to the underlying .swc datafile. To conserve this relationship, but also achieve better computational efficiency, morphological trees in NEAT have the `set_comp_tree()` function. This function internally constructs a coarse-grained tree, referred to as the computational tree, where nodes with identical radii and physiological parameters are grouped in a single cylindrical section. This coarse grained tree is activated with the `as_computational_tree()` context manager. Compute-intensive functionalities activate this computational tree automatically. Note that morphologies can also be resampled arbitrarily, through the `create_new_tree()` function.

### 2.2 NEAT’s ion channels

NEAT provides a simple and lightweight ion channel description format. This format admits ion channels of the Hodgkin-Huxley (HH) type [77], by which we mean that the channel currents are of the form

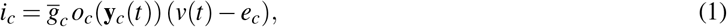

where 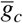 the channel’s maximal conductance density, *e*_*c*_ the reversal potential, and *o*_*c*_ the opening probability as a function of the channel’s state variables **y**_*c*_(*t*). The state variables then evolve according to first-order ODEs that are non-linearly coupled to the local membrane voltage *v*(*t*), which can be specified in two ways: either

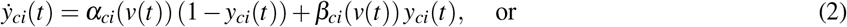

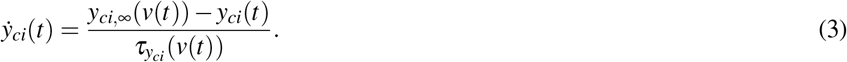

Note that the functions *α*_*ci*_(·), *β*_*ci*_(·) or 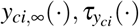 may additionally depend in a user-defined fashion on ion concentrations. Note furthermore that the open probability can be any expression featuring the channel’s state variables.

Implementation-wise, one defines channels as subclasses of `IonChannel`, NEAT’s ion channel base class (Fig 1D, top). Then, it suffices to write a `define()` function, which defines the open probability `p_open` as a channel attribute that is a SymPy expression or a Sympy-readable string [50]. Similarly, the functions *α*_*ci*_(*v*), *β*_*ci*_(*v*) or 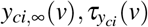 governing state variable evolution are defined as dictionaries with as keys the state variable names and as values the SymPy expressions or SymPy-readable strings. Note that the dictionary keys have to correspond to state variable names featured in the open probability `p_open`. A temperature dependence factor `q10` for the reaction rates can be specified separately, as well as a custom reversal `e` (if not provided, a NEAT default is used based on the ion the channel is permeable to (Na: 50 mV, K: -85.0 mV, and Ca: 50.0 mV). State variable evolution can also depend on a set of ion concentrations, which need to be specified in the `conc` attribute.

NEAT’s `IonChannel` class features functions that provide the necessary quantities to NEAT’s analytical algorithms, as well as code generators that write .mod-files for NEURON and .nestml files for NEST, ensuring that both the analytical computations and the models in various simulators use the exact same underlying channel dynamics. Moreover, groups of NEAT’s ion channels can easily be compiled into ‘models’ for NEURON or NEST, using the ‘neatmodels’ script (Fig 1D, bottom), which then allows identical models to be simulated in both simulators.

### 2.3 The coordinate system of NEAT

To fully specify a location on a branching morphology consisting of cylindrical sections, we need both a node index *n*, and a spatial coordinate *x ∈* [0, *L*_*n*_], which runs from 0 at the proximal end to *L*_*n*_ – the node’s length – at the distal end. For instance, we may specify the membrane voltage at node *n*, position *x* as *v*_*n*_(*x, t*). As the lengths of nodes may vary depending on the morphology specified in the .swc source file, it is not convenient to refer the actual *x*-coordinate in practice. NEAT therefore uses a normalized spatial coordinate 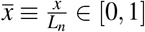. Thus, a location *ξ* in NEAT is always specified as a tuple of node index and normalized x-coordinate, i.e. 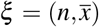. To avoid cluttered subscripts in the mathematical notations, we will adopt the convention that if a quantity, e.g. membrane voltage, features a full location coordinate 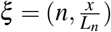, the node index will not be made explicit, i.e. *v*(*ξ, t*) *≡ v*_*n*_(*x, t*).

Locations in NEAT are implemented with NEAT’s `MorphLoc` class, and are always defined with respect to a reference tree (a `MorphTree` or derived class). These locations obey the continuity constrained at the connection points between cylinders: if node *n* is the parent node of *m*, then (*n, x* = 1) *≡* (*m, x* = 0). Furthermore, from the user perspective, these location objects are invariant to the coordinate transformation induced by the switching to the computational tree-context; internally, the `MorphLoc` constructs and stores a coordinate for the computational tree that corresponds to the same location on the original tree.

### 2.4 Model construction

All operations in NEAT revolve around a tree graph structure that stores the morphological and electrical parameters of the neuron model. If the analysis to be performed is limited to the morphological structure of the neuron, a `MorphTree` instance suffices. However, as soon as the electrical behaviour of a cell needs to be studied, physiological parameters need to be added to the neuron model. NEAT implements an intuitive system to build models rapidly. To illustrate the functionalities of NEAT, we have implemented a classical layer 5 pyramidal cell (L5PC) model by Hay et al. [51] (Fig 2A).

**Figure 2.**
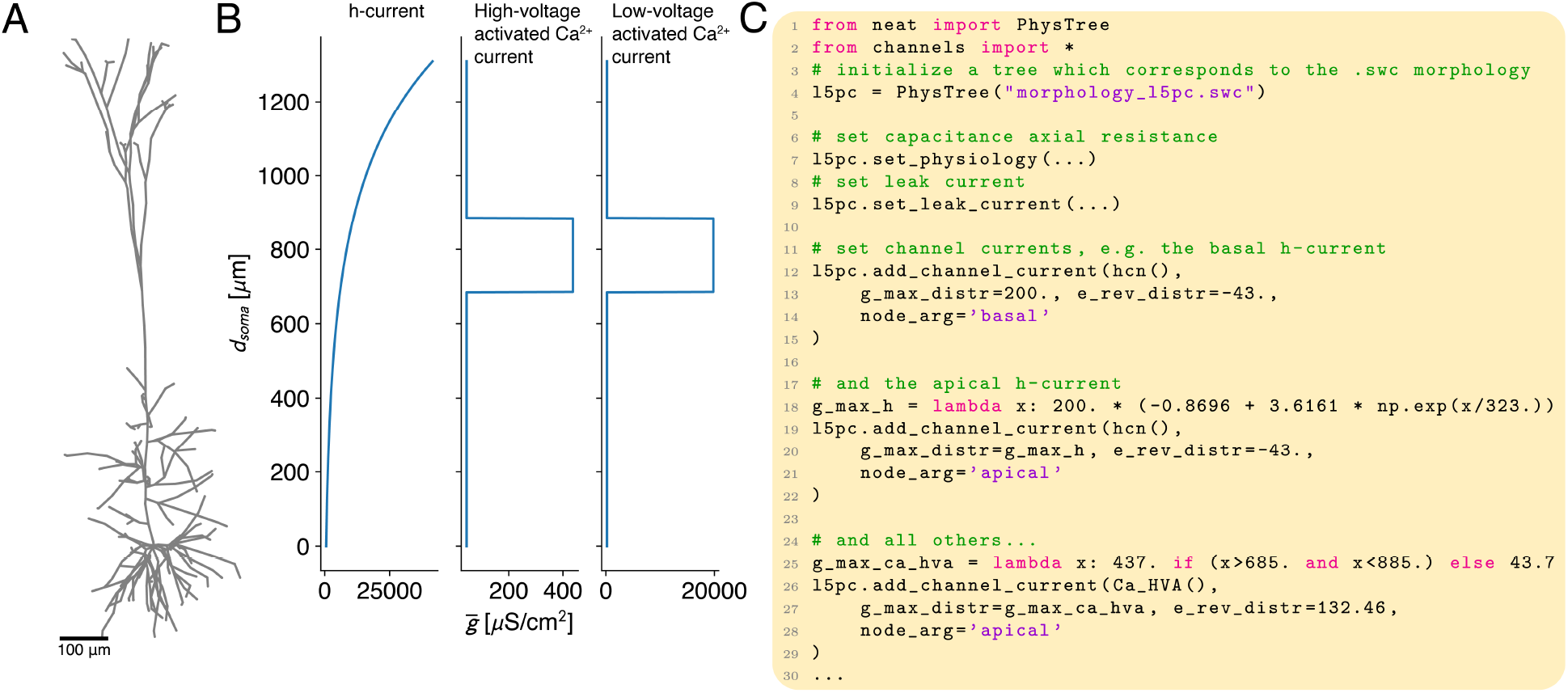
Constructing morphological models in NEAT. **A:** NEAT loads morphologies as .swc files. **B:** Ion channel densities are specified as a function of the distance from the soma (in *µ*m) **C:** Code example of how to load a morphology and how to distribute ion channels. Parameters such as conductance density (`g_max_distr`) and reversal potential (`e_rev_distr`) can be specified as floats, callables where the argument signifies the distance to the soma, or on a per node basis as dictionaries with node indices as keys. Note that the ion channels have to be provided as `neat.IonChannel` objects (here imported from `channels.py`).

First, any morphological tree class instance can be constructed from an ‘.swc’ file, the de-facto standard for describing neuron morphologies (Fig 2A,C, [52]). Constructing the model in this way results in a tree consisting of nodes whose indices correspond to the node indices in the ‘.swc’ datafile (Fig 1C), and whose morphological parameters — cylinder radius, cylinder length, and spatial coordinate of the most distal point (Fig 1C) — contain the values from the underlying datafile.

At this point, the NEAT model only contains morphology information. To turn this into a full, biophysical neuron model, physiological parameters such as membrane capacitance, axial resistance, leak conductance densities and reversals, and ion channel conductance densities and reversals (Fig 1C) have to be set. NEAT’s `PhysTree` and derived tree classes provide convenient functions for these tasks, that provide a great deal of flexibility. They allow these parameters to be set uniformly by passing a single float, as a function of the distance to the soma by passing a callable function, or on a per-node basis by passing a dictionary with node indices as keys (Fig 2B,C). Further, these functions can be applied to specific groups of nodes using the `node_arg` argument.

As many algorithms in NEAT perform analytical calculations on cylinders, ensuring that the number of cylinders in a neuron model is minimal is beneficial for computational efficiency. Often, situations occur where a linear section of dendrite consists of many ‘.swc’ nodes, each with the same radius and physiological parameters. These small cylindrical segments can be grouped into a single cylinder encompassing the whole dendritic segment. However, converting a tree object in this way would distort the morphology, and also result in a mismatch with the underlying ‘.swc’ datafile. For these reasons, morphological trees in NEAT contain a coarse-grained computational tree under the hood. This tree is computed by calling the `set_comp_tree()` function, which should be done *after model construction is complete*. NEAT then provides the `as_computational_tree()` context manager, which activates the computational tree. Note that the associated coordinate transformations are all performed internally by NEAT, so that the user facing interface always expects coordinates that reference the original tree. Note furthermore that modifying *any* physiological or morphological parameters of the neuron model, will automatically result in the deletion of the computational tree, as it can not be guaranteed to be the same after the modification. Note finally, that many of NEAT’s analytical computations will automatically attempt to activate the computational tree context, and raise an error when it has not been set.

### 2.5 Interacting with NEAT trees

NEAT provides a Pythonic way of interacting with its trees. Individual nodes can be accessed by their swc-index (Fig 3A), and NEAT also provides a depth-first iterator for accessing all nodes (Fig 3A). This allows for the selection of specific groups of nodes according to user-defined criteria (Fig 3B).

**Figure 3.**
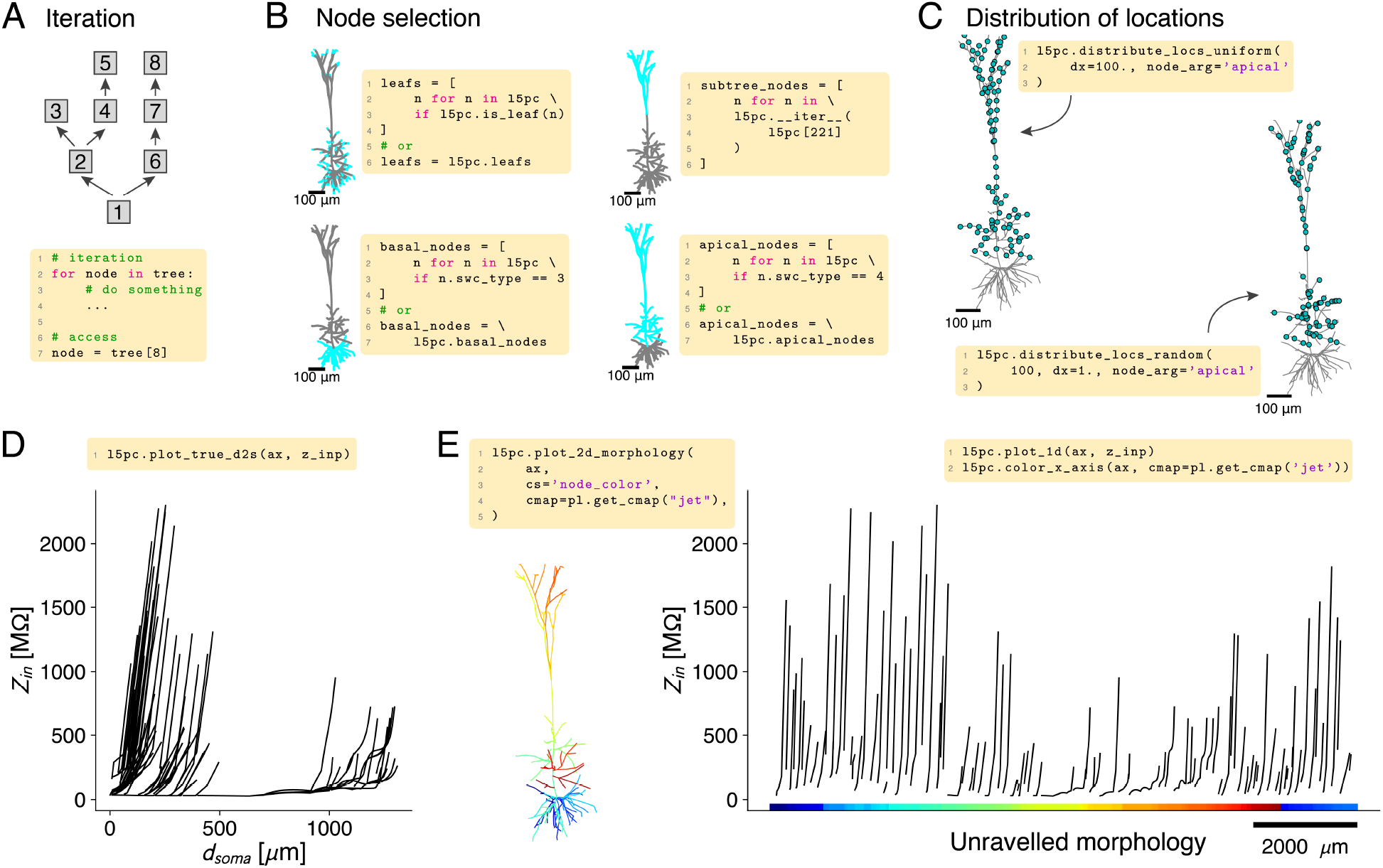
Interacting with morphological models in NEAT to access individual nodes and groups of nodes, to distribute locations, and to plot location-dependent quantities on the morphology. **A:** NEAT trees provide a depth-first iterator, which proceeds to the end of a branch first, and then continues at the nearest bifurcation where there are child nodes that have not yet been traversed (top). Individual nodes can be accessed by their index (bottom). **B**: NEAT allows the flexible selection of groups of nodes through its iterator. Standard groups of nodes, such as the distal tips (leaf nodes, top left), basal nodes (bottom left), or apical nodes (bottom right), can be accessed through convenient attributes. Providing a node to the iterator (top right) starts the iteration from that node, providing access to a subtree of the dendrite. **C:** NEAT’s `MorphTree` and derived classes provide a number of functions to distribute locations, which could represent synaptic input sites or compartment sites, on the tree. These functions provide for instance evenly spaced locations (top left) or randomly distributed locations (bottom right). **D:** Functionality to plot spatial quantities as a function of the distance to the soma, here illustrated with the input resistance of the L5PC model. **E:** Unraveling of the morphology to plot quantities (here illustrated with input resistance as in D). A matching color code between morphological locations (left) and x-axis (right) clarifies the location of each branch.

As described previously, locations on the morphology, which could for instance represent synapse sites or compartment sites, are encoded as a node index and normalized x-coordinate. NEAT provides the `MorphLoc` class for storing such locations, which also stores a reference to the tree on which the location is defined. As such, `MorphLoc` instances are context-aware, and return either original tree coordinates or computational tree coordinates, depending on which tree is active. Nevertheless, all functions that require locations as input also accept simple tuples `(node_index, x)` or dictionaries `{‘node’: node_index, ‘x’, x}`, and interpret them in the coordinate system of the original tree. They are then converted internally to `MorphLoc` instances, so that users are not obliged to instantiate them explicitly. NEAT further provides functionality to distribute locations on the morphology, for instance in a uniformly spaced or random fashion (Fig 3C). These functions return lists of locations, i.e. containing `MorphLoc` instances.

As one of NEAT’s purposes is to analyse dynamics in dendrites, NEAT allows visualization of quantities that depend on the location, such as input or transfer resistances, membrane voltage, concentrations, etc. For instance, such quantities can be visualized as a function of the distance to the soma (Fig 3D). While this provides an accurate view along each branch, it can result in a plot that is hard to parse for complex morphologies that branch frequently. For this reason, NEAT also allows unravelling the morphology on a one-dimensional axis (Fig 3E). In this view, x-positions can be matched to the location on the neuron through the color code of the x-axis. Finally, straightforward recipes to color code quantities that depend on dendritic location directly on the neuron are outlined in the documentation.

## 3 Results

### 3.1 Response kernel calculation

The response kernels, input and transfer impedances, and input and transfer resistances of a neuron model reveal much about how the dendritic tree computes. These are all ways of expressing the Green’s function of the linearised neuron model (see supplement). In particular, response kernels have provided insight into compartmentalisation of dendrites into computational subunits [33], and into the temporal filtering characteristics of dendritic trees [36, 47]. Input and transfer impedances yield insight into frequency-dependent response properties, such as preferred frequencies along the dendritic arborisation [35], and the frequency dependent electrical size of the dendritic tree [53, 54]. Finally, input and transfer resistance have shown to be a strong indicator of compartmentalisation [32, 33, 38], yielding information on spatial interactions between conductances on dendritic trees and the integrative properties of the dendrite. In NEAT, they are furthermore used extensively in the simplification algorithm. Thus, analytical algorithms to compute the Green’s function are the foundation of NEAT.

Each algorithm requires different intermediate quantities to be stored on the tree graph, and therefore is implemented by a separate tree class. NEAT provides three ways of computing the Green’s function: *(i)* using Koch’s algorithm, which assumes the quasi-active approximation (with `GreensTree` / `GreensTreeTime`), *(ii)* using Major’s algorithm to find the separation of variables solution, which only works for passified membranes (see supplement, with `SOVTree`), and *(iii)* through simulations of the full neuron model (with `NeuronSimTree`), where the voltage in response the small, short current pulses is measured (Fig 4A). Koch’s algorithm operates in the frequency domain, but `GreensTreeTime` implements NEAT’s algorithm to convert these impedances to time domain kernels. As Major’s algorithms yields superpositions of exponentials, it applies both to the frequency and time domains. Finally, simulations only yield time domain results (Fig 4B).

**Figure 4.**
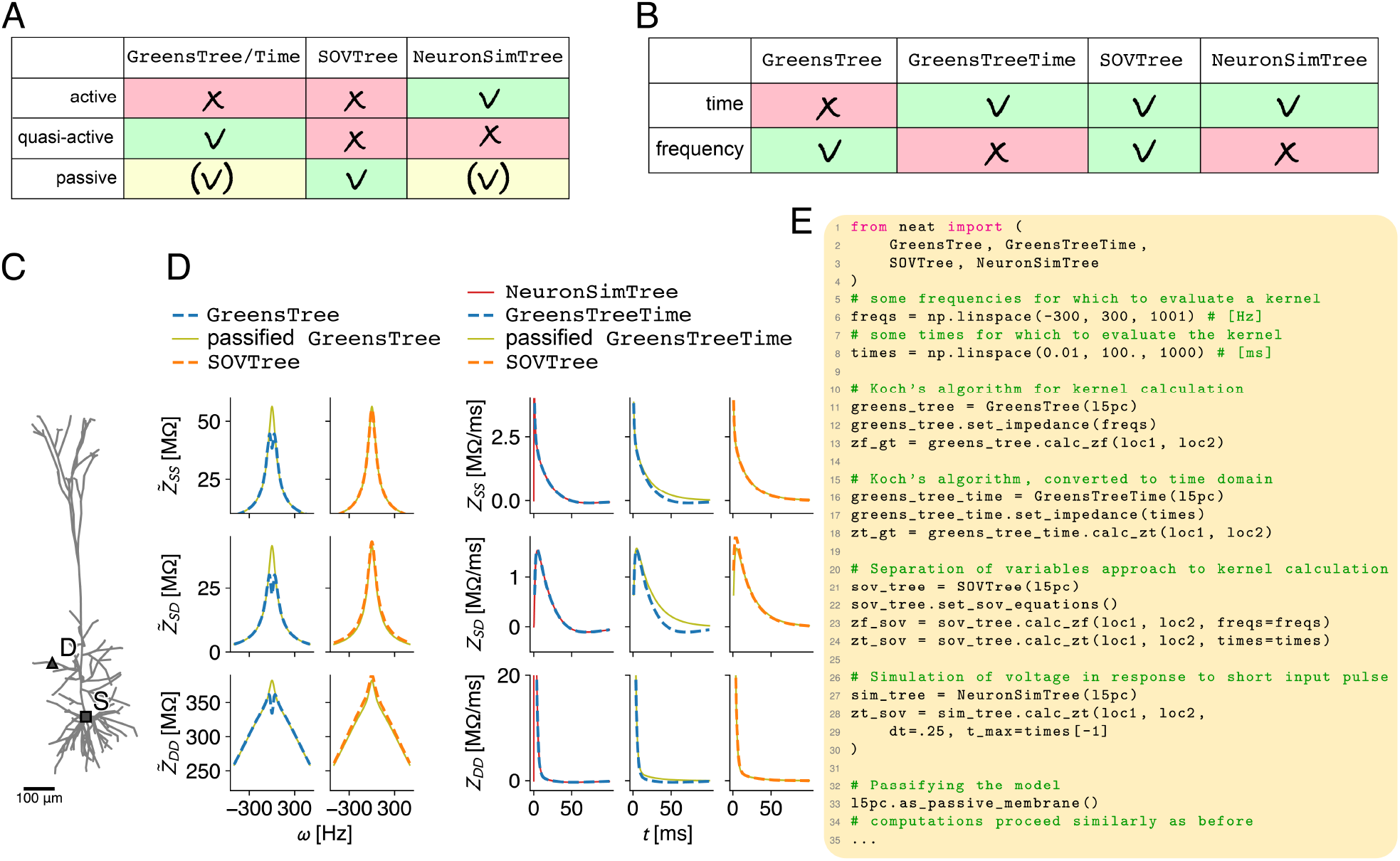
Overview of different input and transfer response kernel calculation methods implemented in NEAT. **A:** Implicit assumption made by each tree class. `GreensTree/Time` and `NeuronSimTree` can be made passive using the `as_passive_membrane()` function. **B:** Overview of whether the kernel calculation method implemented by the tree class computes the response kernels in the time and/or frequency domain. **C:** Exemplar L5 pyramidal cell morphology with a dendritic site (D) and the soma (S). **D:** Input response kernel at the soma (top), transfer response kernel between soma and dendrite (middle), and input response kernel in the dendrite (bottom), computed at site S and D in C. Left, response kernels in the frequency domain. Note that the `GreensTree` by default uses the quasi-active approximation, whereas the `SOVTree` can only use the passive approximation (cf. A). The `GreensTree` can be passified, however, after which the kernels coincide with those computed by the `SOVTree`. Right, response kernels in the time domain. Here, kernels can also be computed by an explicit simulation of the neuron model, by measuring the voltage response following a small and short input current pulse. These kernels coincide with the quasi-active ones computed by the `GreensTreeTime`. **E:** Code example of how to compute the different kernels. With `GreensTree/Time`, all internal impedances (essentially, the boundary conditions for all cylindrical segments, cf. supplement) need to be computed first using the `set_impedance()` function. To compute the kernels as superpositions of exponentials, the transcendental equation needs to be solved first, and then the equations to compute the associated spatial modes. Both steps are performed in the `set_sov_equations()` call (cf. supplement).

To illustrate the different algorithms, we compute the response kernels at a dendritic site, the soma, and between soma and dendrite on the L5PC (Fig 4C), which features a total of seven active ion channels on the apical tree. It can be noted in this regard that impedances computed from the passified tree (i.e. with the explicitly passified `GreensTree` or with the `SOVTree`) are always low-pass filters, and impedances from the quasi-active tree can be band-pass filters (Fig 4D, left). Furthermore, the passified and normal kernels coincide in the high-frequency regime; on short time-scales (below 1 to 10 ms), the dendritic voltage response is dominated by the axial currents, which are captured equally well in the passified model as in the quasi-active one. For this reason, the passified model is an excellent predictor of the compartmentalisation of dendritic trees into computational subunits [33]. In the low-frequency regime - and therefore on longer time-scales (*>* 10 ms) - the response kernels from quasi-active and passive models differ substantially, the former reflecting a preferred frequency characteristic of the dendritic h-current [55, 56]. In the time-domain, the quasi active response kernels from Koch’s algorithm (`GreensTreeTime`) agree well with those computed through simulations (`NeuronSimTree`, Fig 4D, right).

So which algorithm should one choose? All algorithms can be called in few lines of code (Fig 4E). For accuracy and computational efficiency, Koch’s algorithm is the preferred choice. The separation of variables method yields a nice and useful representation as a superposition of exponentials, but is computationally less efficient. Large neurons with many branches furthermore produce many equalising timescales (zeros of the transcendental function) in a similar range, some of which can be missed by the zero finding algorithm. This results in a deterioration of accuracy. The simulation based approach is accurate, but computationally inefficient. It also can not easily be applied to other expansion points than the resting state.

### 3.2 Model reduction

Producing reduced neuron models (Fig 5A) for efficient network simulations is a major use case of NEAT. NEAT’s reduction methodology, as described by Wybo et al. [33], has been extended with the option to incorporate ion concentration dynamics. First, the reduction methodology fits the conductances of the passive model, and then of each ion channel separately, by ensuring that the reduced model approximates the resistance matrix of the full model optimally in the least squares sense (Fig 5B). To generalize well to the full dynamical range, the reduction methodology performs this fit for up to sixteen expansion points per ion channel at the same time. Once this is done, the concentration dynamics can be computed. NEAT’s aim is to achieve the same ion concentration in the reduced compartment as in the full model at the corresponding site. To achieve this in practice, it has proved sufficient to rescale the total ionic current by a factor computed from the fit of the ion channels that carry the ion under consideration (see supplement). The capacitances are then computed to reproduce the largest eigenmode of the full model, computed from the separation of variables expansion, or by requiring the same membrane time scale as at corresponding locations in the full model. Sanity of either of these fits can be checked by comparing response kernels in the full and reduced models (Fig 5C), for which `CompartmentFitter` provides the `plot_kernels()` functions. Finally, the reversal potentials of the leak current are used as a compartment-specific free parameter that fits the equilibrium potentials in the reduction to those in the original model at the compartment locations.

**Figure 5.**
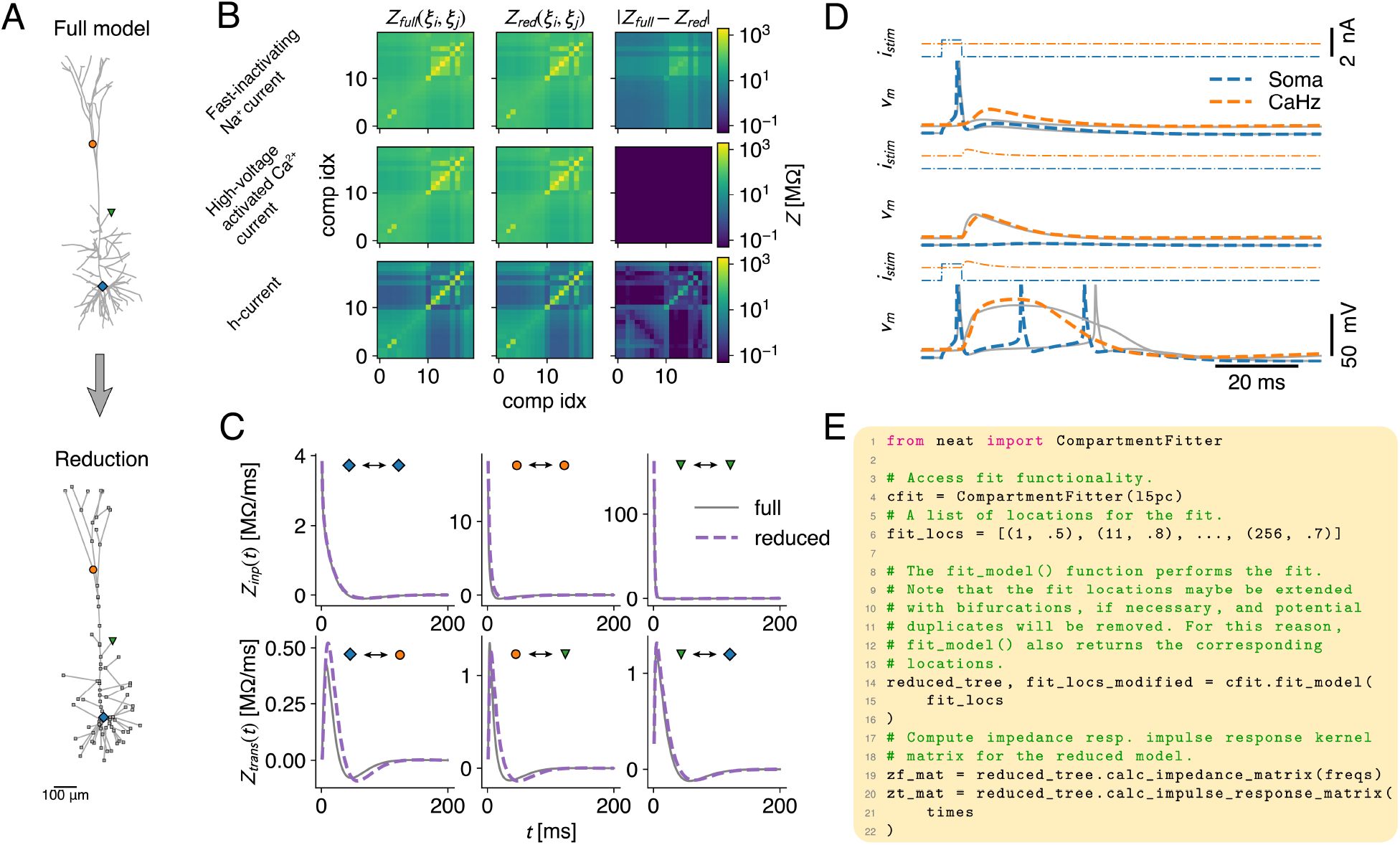
Reducing morphological neuron models with NEAT. **A:** Morphogical models – consisting of cylindrical sections – can be converted to simplified compartment models, where the compartments represent user-defined locations on the morpholgy. **B:** Reductions with NEAT fit the conductance parameters in each compartment to reproduce resistance matrices, i.e. the matrix *Z* where 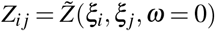. For each ion channel separately, multiple resistance matrices are compute at different activation levels to ensure generalization to the full non-linear dynamics. **C:** Reductions with NEAT fit the capacitance parameter in each compartment either by requiring that the largest eigenmode between full and reduced models matches, or by ensuring that the local membrane timescale is the same as in the full model. Both strategies have been found to reproduce the transfer and input response kernels in the full model (symbols match locations in A). **D:** Simulation of the reduced model (dashed, coloured lines) undergoing the BAC-firing protocol, compared to the full model (solid, grey lines). **E:** Code example demonstrating the simplification workflow. NEAT uses the `CompartmentFitter` class to create reductions based on provided sets of fit locations. The reductions are returned as `CompartmentTree` class instances.

The L5PC model by Hay et al. [51] that was optimised with an evolutionary algorithm to reproduce backpropagation-activated calcium (BAC) firing, constitutes a challenging test case for the fit methodology. This L5PC model contains fast dendritic sodium channels, slow calcium channels with a high conductance density within the Ca-hotzone (cf. Fig 2B,C), and an h-current increasing exponentially with distance from soma. We ran an identical BAC firing protocol in the original (Fig 5A, top) and reduced model ((Fig 5A, bottom) and found qualitatively similar dynamics.

To construct a simplified model in NEAT, it suffices to create a `CompartmentFitter`, specify a list of locations on the morphology where the compartments should placed, so that they reproduce the dynamics of the full model at those sites, and then call the `fit_model()` function (Fig 5E). For technical reasons, this function may append other compartments locations, such as bifurcations that are in between compartment sites [31]. As a consequence, the number of compartments in the reduced `CompartmentTree` may be larger than the number of locations provided to `fit_model()`. To have a reference to all locations in the reduction, the `fit_model()` function returns the extended list of compartments together with the reduced `CompartmentTree`. It should be noted that that the order of the original locations provided to `fit_model()` will not be scrambled; if *n* locations are provided, the first *n* entries in the returned location list will correspond to the entries in the original location list. Furthermore, each node in the reduced `CompartmentTree` stores the index of the location in the returned location list that it references, so that they can always be mapped back to the original locations.

### 3.3 Simulating full and reduced models in NEURON and NEST

NEAT is not a simulator itself, but a model analysis and reduction tool that exports models to simulators. NEAT can export morphological models to NEURON as connected cylindrical sections, so that NEURON will determine the discretisation in compartments internally. NEAT can also export compartmental models to NEURON and NEST. NEAT has two ways of constructing compartmental models for simulation (Fig 6A): (i) through its reduction methodology, and *(ii)* through its implementation of the second order finite difference approximation (cf. Fig S1). In all these cases, the `IonChannel`s that are included in the model need to be compiled first. For this purpose, NEAT provides the `neatmodels` script, which can be called from the command line (Fig 1D). Additional mechanisms, such as synaptic receptors, can be included with this script in the model compilation as options. When starting a NEURON or NEST simulation, the compiled model must be loaded first, using the `load_neuron_model()` or `load_nest_model()` functions provided in NEAT, so that it is visible to NEURON or NEST (Fig 6B). Then, the tree classes `NeuronSimTree` on the one hand, and `NeuronCompartmentTree` and `NestCompartmentTree` on the other hand, construct respectively the morphological model in NEURON, or the compartmental model in NEURON or NEST (Fig 6B).

**Figure 6.**
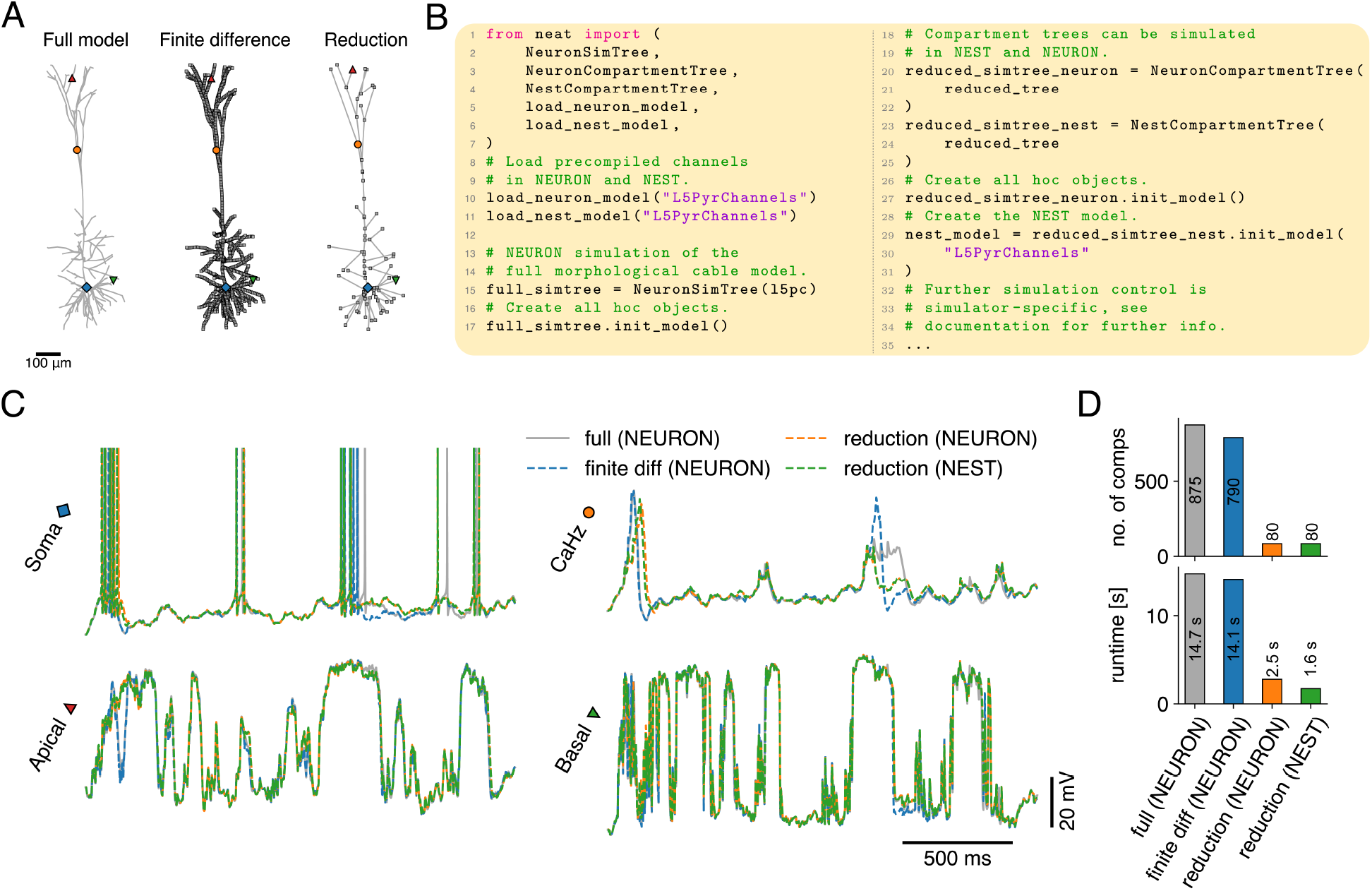
Using NEAT to create simulations in NEURON and NEST. **A:** NEAT can be used to export morphological models – consisting of cylindrical sections (left, full model) – to NEURON, and compartmental models to NEURON and NEST. NEAT provides two ways of obtaining compartmental models from full morphological models: either through the reduction approach (right, reduction) or through NEAT’s finite difference approximation (middle, finite difference). **B:** Code example of exporting models to simulators. NEURON simulations of morphological models – where NEURON internally determines the discretisation (i.e. the conversion to a compartmental description) – can be constructed with a `NeuronSimTree`, whereas NEST resp. NEURON implementations of compartmental models can be constructed with `NestCompartmentTree` resp. `NeuronCompartmentTree`. Further simulation control is simulator specific, as each simulator has its own API. **C:** Comparison of voltage dynamics under NEURON’s discretization (grey), NEAT’s finite difference discretization simulated with NEURON (blue), NEAT’s reduction simulated with NEURON (orange), and NEAT’s reduction simulated with NEST (green). Locations as indicated in A. **D:** Number of compartments in the resulting model (top) and the associated runtime for a simulation of 2 s (bottom), simulated on a MacBook M3 Max (NEURON version 8.2.4 and NEST version 3.8.0, both installed following standard procedure as outlined in the respective documentations).

To compare the compartmentalization methods, reductions and simulators, I have constructed simulations where identical Poisson input trains to glutamatergic (featuring AMPA and NMDA receptors) and GABAergic synapses impinged at equivalent locations (Fig 6C). Generally, I find good agreement between responses in the different models, especially in apical and basal subunits. The precise waveform of the Ca^2+^ spike can differ for different complexity levels and even between NEURON and NEAT’s discretization schemes (cf. supplement) on a case by case basis, highlighting the challenging nature of accurately representing highly non-linear spatio-temporal dynamics.

Nevertheless, timing of the associated action potential bursts matched for all cases. As model runtimes scale approximately linearly with the number of compartments, the reduced model – which features approximately ten times fewer compartments than the full models (Fig 6D, top), has a similarly reduced runtime in NEURON (Fig 6D, bottom). Interestingly, the NEST implementation achieves a speedup by a factor of almost two over the equivalent NEURON model.

## 4 Availability and future directions

NEAT can be installed through pip (https://pypi.org/project/nest-neat/) or from the source repository (https://github.com/nest/NEAT). Further documentation on NEAT can be found at https://neatdend.readthedocs.io/en/latest/reference/index.html. The full code to reproduce the figures from this manuscript can be found at https://jugit.fz-juelich.de/w.wybo/NEATPaper.

Classical analytical methods to compute voltage responses in the dendritic tree can reveal a great deal about dendritic computation, even in the presence of non-linear, voltage-dependent currents. Here, we have highlighted the reason for this: at a short time-scale, dendritic voltage dynamics are dominated by axial currents, which are passive. Therefore, active transmembrane currents can be seen as being integrated by a scaffolding provided by the passive system.

This provides an improved understanding of spatiotemporal interactions between dendritic excitation and inhibition [33, 38, 57, 58]. Further, it reveals that different loci on the dendritic tree respond preferentially to different input frequencies [35, 59–64]. However, due to the absence of modern implementations, usage of these algorithms to understand dendritic computation has remained limited, with authors often relying on simplistic morphologies while investigating analytical solutions (e.g. the ‘ball-and-stick’ model, with a linear dendrite) [65–67]. Such simple morphologies, however, do not produce the same input resistance profiles from soma to tip as real morphologies, where dendritic segments branch and taper towards the tips, and should therefore be treated with some care. By implementing these algorithms for arbitrary morphologies, NEAT fills this gap in the software toolset of neuroscience.

A powerful usage of these classical analytical methods, in particular Koch’s method to compute quasi-active voltage responses, is to compute large numbers of resistance matrices of the full model, for many expansion points in parrallel. This can not be achieved efficiently through simulations, and, as demonstrated by our prior work [31], can be used to constrain the conductance parameters of reduced models. The resulting reductions faithfully approximate full models, but require much less time to simulate, as the simulation cost scales linearly with the number of compartments. NEAT implements functionality to export these reduced models to NEURON [1] and NEST [2, 68, 69]. In the future, NEAT plans to support exporting models to other simulators, such as Brian [3] for models that are easy to adapt, Arbor [12,13] for biophysically detailed models, or Jaxley [70] to fine-tune parameters.

NEAT exports models to two of the most commonly used simulators in neuroscience, and reads in the most commonly used morphology description format. As such, NEAT is already well embedded in the neuroscientific toolchains. However, NEAT defines its own ion channel format, that is for now also limited to Hodgkin-Huxley type ion channels, as NEAT performs many analytical computations using the ion channels. A more general ion channel system is planned for a future version. Nevertheless, the definition of ion channels is compact and easy to specify. Once they are implemented, they can be compiled into NEURON’s ‘.mod’-files [1] or NESTML models [5, 68, 71], which are in turn compiled into efficient simulation codes by these software packages. All of these steps can be performed through a single terminal command (Fig 1D).

Reduced models of dendritic computation can be grouped broadly speaking in two categories: *(i)* top-down, where the model is built up to capture a particular dendritic computation, in a manner that is as simple as possible [21–25], and *(ii)* bottom-up, were starting from a full morphology, a mathematical simplification procedure is applied to arrive at a simpler model [27, 28, 30, 48]. NEAT’s simplification algorithm can be employed in both cases [25, 31], as it provides the freedom to focus on specific computations by retaining appropriate sets of locations on the morphology, while accurately capturing both intra-dendritic and somato-dendritic interactions. In modern approaches, it is often desirable to capture multiple dendritic computations in a realistic fashion [23]. NEAT was used, for instance, to show that a two-compartment adaptive exponential integrate-and-fire model that exhibits BAC-firing, can be extended with NEAT to have further apical and basal compartments that support NMDA spikes [25].

Taken together, NEAT fills a gap in the neuroscientific software toolset by on the one hand allowing the study of single neuron properties in great detail, through its analytical algorithms and model export to NEURON functionality, and on the other hand by its functionality to embed simplified versions of the *same* neuron model in large-scale networks, through its export to NEST functionality. As such, NEAT helps to bridge a long-lasting divide in computational neuroscience, where research was either focused on one or a few highly detailed models, or on large networks of highly simplified point-neurons.

## Acknowledgements

This work was supported by Helmholtz Association’s project-oriented funding programme (PoF 2, Topic 3). I gratefully acknowledge computing time on the supercomputer JURECA [72] at Forschungszentrum Jülich under grant jinm60. Further, I would like to extend my gratitude towards Jessica Mitchell for support with NEAT’s documentations website, towards Dennis Terhorst for providing software development support and coordination, and towards Hermann Cuntz and Alexander Bird for providing invaluable comments on this manuscript. Finally, I would like to thank Abigail Morrison, and by extension the whole board of directors of the NEST Initiative, for believing in this project.

## 5 Supplementary information

### 5.1 Mathematical formulation of morphological neuron models

#### 5.1.1 Neurons as systems of coupled cables

In the morphological tree classes in NEAT, nodes are taken to represent cylindrical sections of neurite, which are connected at their proximal end to the cylindrical section associated with their parent node, whereas at their distal end they connect to the cylindrical sections associated with their child nodes. On such a cylindrical section associated with node *n*, membrane voltage is modeled by the cable equation [73], which was first applied to dendrites by Rall [74–76]. With the inclusion of active channels and synaptic inputs, this equation has the following form:

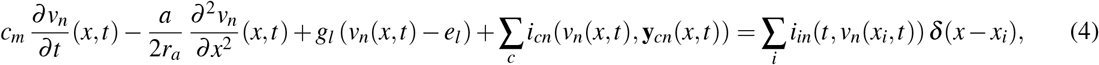

where *c*_*m*_, *r*_*a*_, *g*_*l*_, and *e*_*l*_ denote, respectively, the membrane capacitance, the the axial resistance, the leak conductance and the leak reversal, *a* denotes the radius of the dendritic branch, *i*_*c*_ the current contribution of a channel type *c* and *i*_*i*_ the a-priori arbitrary input current at location *x*_*i*_. The ion channel current *i*_*c*_ can depend non-linearly on the voltage and a number of state-variables **y**_*cn*_. Note that the parameters are all specified on a per-node basis, but for notational clarity the node index was not made explicit. At the moment, NEAT only accommodates channel currents of the Hodgkin-Huxley type [77]:

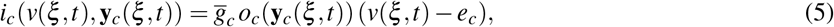

so that the channel current depends on a maximal conductance density 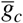, a driving force (*v*(*x, t*) − *e*_*c*_) – with *e*_*c*_ the channel’s reversal potential – and *o*_*c*_(**y**_*c*_(*ξ, t*)) the channel’s open probability, which depends on a number of state variables **y**_*c*_(*ξ, t*) = (*y*_*c*1_(*ξ, t*), …, *y*_*cK*_(*ξ, t*)) that evolve according to

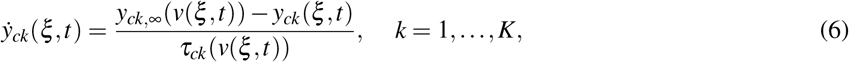

with *τ*_*ck*_(*v*) and *y*_*ck*,∞_(*v*) functions that depend on the channel type. In NEAT, all parameters are stored on a per-node basis. Hence, each cylindrical section has spatially uniform parameters, but they can have different values for each node. Note furthermore that NEAT only considers the left-hand side of equation (4) in its analysis tools. A-priori arbitrary sets of inputs can be added in the models that are exported for simulation.

Voltage dynamics are constrained by boundary conditions [73]). When tree nodes are leafs (i.e. they have no child nodes, and represent the most distal segments of neurite), we assume a sealed-end boundary condition (no longitudinal current flow:

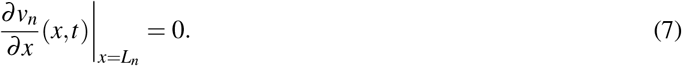

Where two or more cylinders join together, the boundary conditions are given by requiring the equality of the membrane potential:

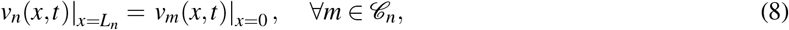

and conservation of longitudinal current flow:

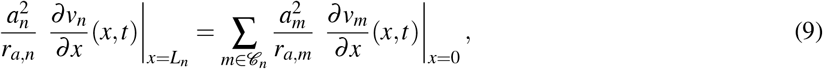

where 𝒞_*n*_ denotes the set of child nodes of node *n*. Finally, at the soma, we apply the lumped-soma boundary condition. To this purpose, the root node of the tree, which in NEAT is assumed to implement a spherical soma of radius *a*_*s*_, is modeled electrically as a single compartment with voltage *v*_*s*_. The boundary conditions then stipulate equality of the membrane potential:

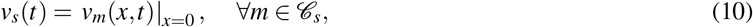

and require that the total current flow through the somatic membrane must equal the axial current flow through the branches emanating from the soma:

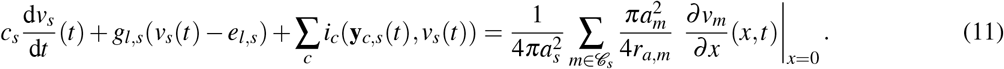

Here, somatic voltage and parameters are denoted with the subscript *s*.

#### 5.1.2 The quasi-active approximation for computing linearized neural dynamics

The system of equations described above can be linearized around an a-priori arbitrary expansion point *v*(*ξ, t*) = *v*_0_(*ξ*) + *δv*(*ξ, t*) for voltage and **y**_*c*_(*ξ, t*) = **y**_*c*,0_(*ξ*) + *δ* **y**_*c*_(*ξ, t*) for ion channel state variables [78]. In NEAT, this expansion point can be specified on a per-node basis, and hence is constant for each cylindrical section; NEAT implicitely assumes the approximation *v*_0_(*ξ*) *≡ v*_0,*n*_(*x*) *≈ v*_0*n*_ and **y**_*c*,0_(*ξ*) *≡* **y**_*c*,0,*n*_(*x*) *≈* **y**_*c*,0,*n*_. Linearizing ion channel currents then yields a quasi-active description [54] of the neuron model:

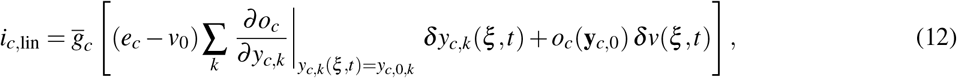

with

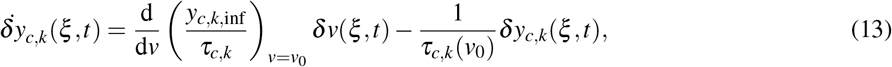

where all derivatives, as well as *τ*_*c, j*_, are evaluated at the expansion point of the state variables. If the expansion point given by *v*_0_(*ξ*) and **y**_*c*,0_(*ξ*) is stable, then *δv*(*ξ, t*) and *δ* **y**_*c*_(*ξ, t*) will model the responses to transient input perturbations that are small enough so that the linear approximation around the expansion point is valid. A natural special case of this is the situation where the expansion point is the equilibrium of the neuron in the absence of external inputs. The linearization then models responses to small inputs. This quasi-active approximation is assumed by the `GreensTree` and the `GreensTreeTime` in their implementation of the Green’s function calculation.

#### 5.1.3 The membrane leak approximation for computing passified neural dynamics

A further approximation of (4) can be obtained by replacing the dynamic channel open probabilities *o*(**y**_*c,n*_(*x, t*)) with their value at the expansion point *v*_0,*n*_, **y**_*c*,0,*n*_, so that the channel conductance 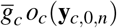 is a constant which is added to the leak:

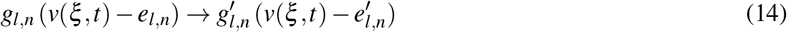

where

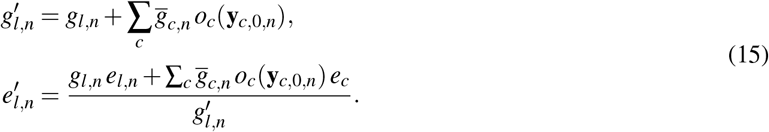

In doing so, the neuron model is converted to a passive one. Despite the strong reduction, this approximation still provides useful information. In particular, the longitudinal spread of the voltage is dominated by the axial current, and therefore morphological properties such as the radii of the cylinders. By consequence, this ‘passified’ neuron model is remarkably accurate in reproducing the effective compartmentalization [33] of the original model (cf. Fig 4). Furthermore, NEAT’s `SOVTree`, which implements the seperation-of-variables solution, assumes this description, as this solution has only been derived for passive neuron models. Finally, reducing certain sections of the dendritic tree to the passive approximation may yield sizeable efficiency gains in simulations, and could be justified with respect to certain computations. For instance, local subunits arising through the non-linearity of the NMDA-receptor can be reproduced in such an approximation [33]. To passify models or certain groups of nodes, NEAT’s `PhysTree` and derived classes provide the `as_passive_membrane()` function.

### 5.2 The Green’s function of a morphological neuron model

In essence, the Green’s function *Z*(*ξ, ξ*_*i*_; *t*) is the solution of the system of linearized, coupled cable equations to a delta pulse input current *δ*_*nm*_*δ* (*x* −*x*_*i*_)*δ* (*t*) occurring at a certain site 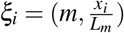 on the neuron at time *t* = 0 [54, 73]. From this, the solution to a general input current perturbation 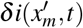 can be constructed through convolution:

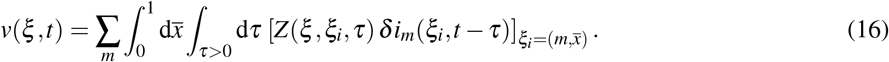

Here the sum runs over all nodes *m*, and the spatial integral integrates over all the associated cylinders. In nearly all practical use cases, the inputs occur at a discrete set of point-like locations in space (e.g. synaptic receptor sites or electrode current injection sites), which will referred to as input sites {*ξ*_*i*_} for notational simplicity. Mathematically, the input is thus a combination of spatial delta functions, allowing us to reduce the compound spatial sum and integral in (16) to a simple sum, i.e.

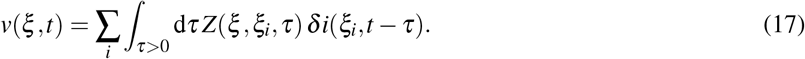

The significance of the Green’s function is that it encapsulates all effects induced by the morphology in a single function, and therefore contains much information about the neuron’s response properties. For instance, input or transfer resistances, measured in response to e.g. DC current steps, are given by the integral *Z*(*ξ, ξ* ^*′*^) = ∫ _*τ>*0_ d*τ Z*(*ξ, ξ* ^*′*^, *τ*). The Fourier transform of the Green’s function, 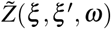, yields the input or transfer impedances, i.e. the voltage response amplitude and phase at a site *ξ* in response to a sinusoidal input current at site *ξ* ^*′*^. Furthermore, *Z*(*ξ, ξ* ^*′*^, *t*) is also known as the impulse response kernel, and contains information on the response time-scales induced by the neuronal morphology. Finally, constructing simplified models that approximate parts of *Z*(*ξ, ξ* ^*′*^, *τ*) as well as possible has proven to be a powerful method of reducing complex neuron models [31].

#### 5.2.1 Computing the Green’s function in the frequency domain: input and transfer impedances

By Fourier transforming (17) to the frequency domain, the temporal convolution becomes a multiplication:

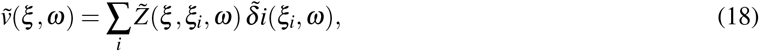

which allows for the straightforward computation of the voltage response amplitude and phase in response to specific input currents. By analysing the impedance 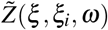, it can be shown that while passive models are always low-pass filters of the input current, ion channels can cause neurons to become band-pass filters, preferentially responding to certain frequency bands [35, 59–64]. Furthermore, the specific preferred frequency depends on the location on the morphology [35]. Koch’s algorithm [39], based on graphical rules derived by Butz and Cowan [79], computes these impedances exactly on tree graphs. If the graph were to contain loops, which might be induced if the impedance needs to be calculated for multiple neurons connected by gap junctions, Abbott’s sum-over-trips approach provides an approximate solution [41, 80, 81].

NEAT implements Koch’s algorithm through the `GreensTree` class, which is briefly sketched here. First, it should be noted that Fourier transforming the linearized neuron model yields a coupled system of 2nd-order ODEs, as temporal derivatives are replace by multiplication with *iω* – where *i* is the imaginary unit. For the linearized cable equation, we obtain

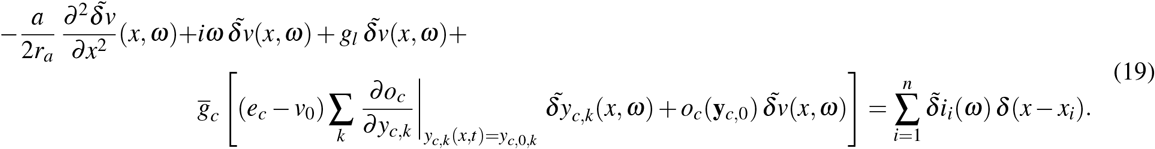

The Fourier transformed ion channel state variables can then be eliminated from this equation, by rearranging the Fourier transforms of their quasi-active evolution equations (13):

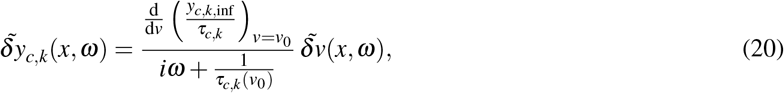

yielding

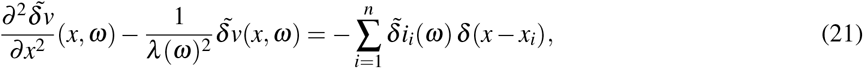

where the frequency-dependent length constant *λ* (*ω*) is given by:

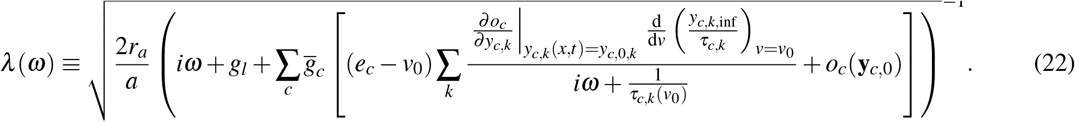

In the remainder of this section we will drop the dependence on *ω* for notational clarity.

It can be seen that cosh(*x/λ*) and sinh(*x/λ*) are two linearly independent fundamental solutions of the homogeneous version of (21), i.e. where the input (right-hand side) is zero. Under homogeneous boundary conditions

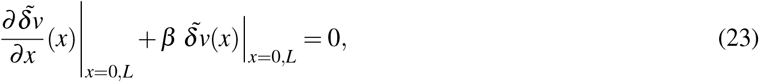

the solution to a delta-pulse input at *x*_*i*_ is:

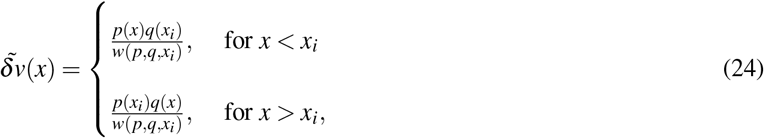

with *p* resp. *q* linear combinations of the fundamental solutions that satisfy the homogeous boundary condition resp. at *x* = 0 and at *x* = *L*, and *w*(*p, q, x*) = *p*(*x*) *q*^*′*^(*x*) −*q*(*x*) *p*^*′*^(*x*) the Wronskian of the solutions [82].

The issue is that at the junctions between coupled cylinders, the boundary conditions (8) and (9) are not homogenuous (and neither is the lumped-soma boundary condition). However, starting from the leaf cylinders, where it can be seen that the sealed-end condition (7) is homogenuous, homogenuous boundary conditions can be constructed for the whole tree. Let 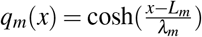 be the solution on such a leaf cylinder *m* that satisfies (7) on the sealed end. Every non-trivial solution on this cylinder, in the absence of direct input to the cylinder, will be of the form *α*_*m*_ *q*_*m*_(*x*) for some factor *α*_*m*_. At the junction with the parent cylinder *n*, we get that 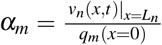 from (8), which leads to a constraint on the derivative:

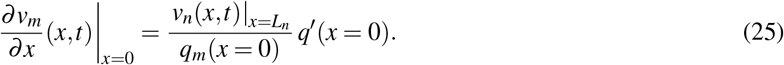

Substituting this in (9) then yields:

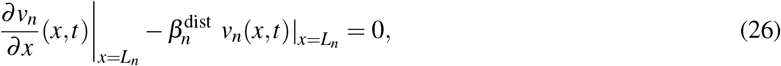

where 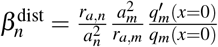. It can be seen that this is precisely the homogeneous boundary condition we were after. In the parent cylinder *n*, we can then again construct a homogeneous solution *q*_*n*_(*x*) from the fundamental solutions cosh(*x/λ*_*n*_) and sinh(*x/λ*_*n*_), that satisfies the constructed boundary condition (26). Note that this scheme can readily be extended to a junction where the parent cylinder has more than one child node, where we obtain for 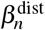:

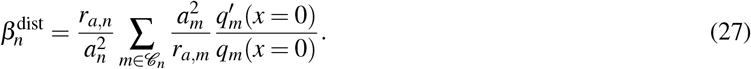

This scheme can then be applied recursively through the tree graph. When the recursion arrives at the soma, working out the lumped soma boundary condition in the same way results in a homogenuous boundary condition (23) on the proximal end of each of the child cylinders of the soma. Continuing the recursive scheme, but now towards the leafs, then yields homogenuous boundary conditions at the proximal ends for each cylinder (for which we call the associated factors 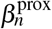. When the `GreensTree.set_impedance()` function is called, NEAT computes the factors 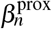 and 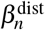 for each node *n* through this recursive algorithm.

Finally, to compute the Green’s function in response to a delta pulse at location *x*_*i*_ on node *n*, NEAT uses (24), where *p*_*n*_(*x*) and *q*_*n*_(*x*) are computed from the homogenuous boundary conditions (23) on proximal resp. distal ends, using the stored factors 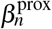 resp. 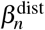. On the child cylinders *m ∈* 𝒞_n_, the solution is then of the form *α*_*m*_ *q* _*m*_ (*x*), with *q*_*m*_ (*x*) computed from the fundamental solutions by satisfying the distal homogenuous boundary condition (i.e. computed from 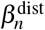). The continuity of voltage condition (8) at the junction between cylinder *n* and cylinder *m* is then used to compute *α*_*m*_, and this scheme is repeated recursively until the dendritic tips are reached. Similarly, on the proximal end – in the parent cylinder *k* we use the solution *α*_*k*_ *p*_*k*_(*x*) which satisfied the proximal homogenuous boundary condition (i.e. computed from 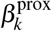), where the prefactor *α*_*k*_ is again computed from the continuity of voltage condition, now at the junction between cylinders *k* and *n*. This scheme is then repeated recursively until the soma is reached. Note that in practice, the Green’s function is always evaluated between a discrete set of locations, e.g. through calling `calc_zf()` or `calc_impedance_matrix()`, and the recursion will only be applied to the direct path between the input location and the target location(s).

#### 5.2.2 Inverse Fourier transform algorithms to compute the Green’s function in the time domain

While the `GreensTree` computes the impedances exactly in the frequency domain, time-domain impulse response kernels are obtained in NEAT with `GreensTreeTime`, which implements algorithms that compute the inverse Fourier transform

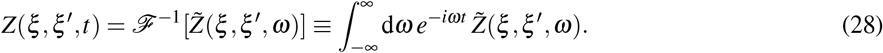

Computing this quadrature accurately is not a straightforward problem: on the one hand, these kernels need to be accurate on submillisecond times-scales, to capture voltage transfer between nearby locations, while on the other hand, they need to be accurate on long time-scales (10-100 ms), to capture the voltage decay through the membrane. This means that the quadrature must be computed precisely for small frequencies (long time-scale accuracy), but that it also must incorporate very large frequencies (for short time-scale accuracy). To that purpose, NEAT by default defines an integration grid {*ω*_*i*_} that is composed of 100 equispaced frequency values between -10 and 10 Hz, and a log-spaced frequency grid containing 200 values from 10 to 10^7^Hz (and similarly from −10^7^ to −10 Hz. Note that with this grid, 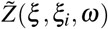 was empirically found to always change smoothly between grid points. However, computing the quadrature (28) naively would still be highly inaccurate, as *e*^−*iωt*^ might oscillate strongly as a function of *t*, with a period smaller than the distance between integration grid points. To resolve this issue, NEAT linearly interpolates 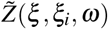 between grid points:

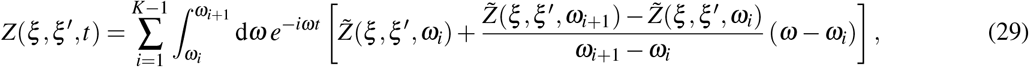

with *K* the number of grid points, and computes the resulting integrals 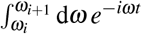 and 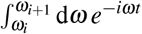 exactly. Finally, these integrals are precomputed for each desired *t*-value at which the impulse response kernel is to be evaluated, and are arranged in a matrix so that the whole quadrature is recast as a matrix-vector product with the vector 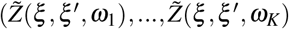.

The quadruature-based approach works well for transfer kernels, where *ξ*≠*ξ* ^*′*^. This is the case because transfer kernels are continuous everywhere (in particular at *t* = 0, as they are identically zero for *t <* 0 (implementing causality), are zero at *t* = 0 (it takes a non-zero amount of time for a current injection at a site *ξ* ^*′*^ to cause a voltage perturbation at site *ξ*≠*ξ* ^*′*^, and then reach nonzero values for *t >* 0. This means that the frequency domain transfer impedance decays to zero for *ω →* ∞, and in particular that 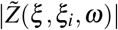 is vanishingly small at the edges of the quadrature grid (i.e. *ω* = −10^7^ and 10^7^Hz). For input kernels however, where *ξ* = *ξ* ^*′*^, *Z*(*ξ, ξ, t*) jumps discontinuously to a non-zero value at *t* = 0 (a current injection at site *ξ* will instantaneously cause a voltage perturbation at that site), the associated input impedance will contain all frequencies, and converge to a non-zero value for *ω →* ∞. Concretely, this means that any quadrature approach will create an artificial cutoff frequency, in turn inducing spurious oscillations in the time domain kernel. To resolve this issue, it can be noted that input kernels decay from their initial value. This decay can be approximated accurately as a superposition of exponentials:

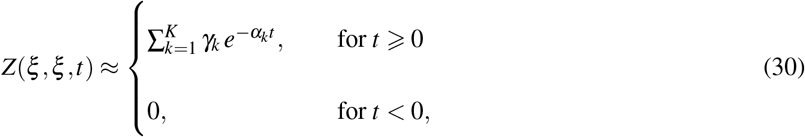

with Re(*α*_*k*_) *>* 0. By default, NEAT sets *K* = 20, as this was empirically found to be a good compromise between speed and accuracy. In the frequency domain, such a representation is of the form:

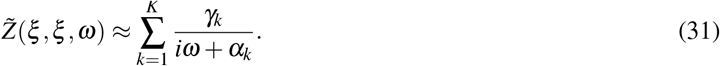

NEAT implements the vector fitting algorithm [83, 84] to approximate input impedances in the frequency domain in this way. For the desired *t*-values, the input kernels are then computed according to (30).

#### 5.2.3 The Green’s function as a superposition of exponentials: the separation of variables solution

The Green’s function can also be computed directly as a superposition of exponentials

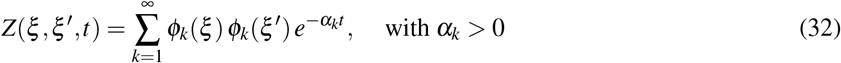

through the separation of variables expansion, which yields an infinite series of response time-scales 1*/α*_*k*_ and associated spatial profiles *ϕ*_*k*_(*ξ*) [43–47]. This formulation has as advantage that the voltage response can be decomposed in a number of distinct modes, each with an associated timescale, which can be seen from substituting (32) in (17):

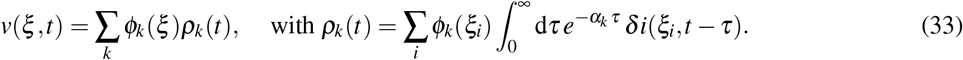

So far, the separation of variables expansion has only been formulated for passive neuron models. In NEAT, this means that it is automatically applied to the passified version (cf. (14), (15)) of (4). First, solutions to the homogeneous problem are computed. On a single cylinder, the passified cable equation has the form (4):

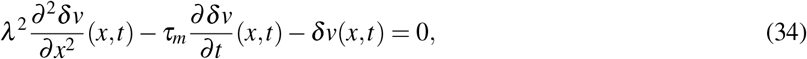

with 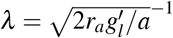 the length constant and 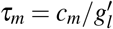 the membrane time constant. In the separation of variables expansion, solutions to (34) are proposed of the form:

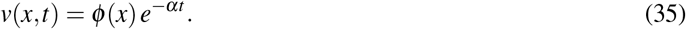

Substituting this in (34) yields:

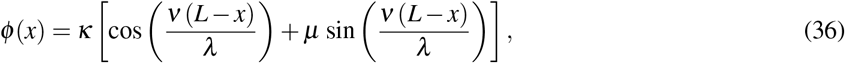

where *κ* and *µ* are parameters to be determined from the boundary conditions, and 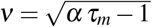. Major’s algorithm then determines the parameters *µ* in a recursive procedure, starting from the distal tips. Here, the sealed end condition holds, leading to *µ* = 0. Applying the continuity of voltage condition (8) at junctions between nodes then yields:

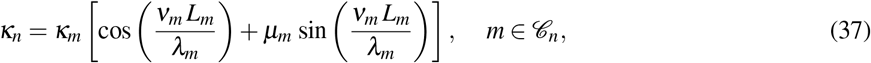

where we have reintroduced the indices *n* for parent cylinder en *m ∈ C*_*n*_ for the child cylinders. The conservation of current condition (9) stipulates:

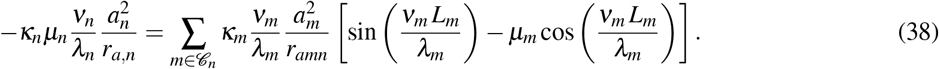

From this, the ratio *κ*_*m*_*/κ*_*n*_ is eliminated, to find a recursive relation that solves for *µ*_*n*_ throughout the tree:

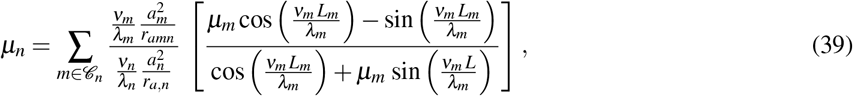

where we note that these analytical expressions depend on the yet unknown constant *α*, since *ν* depends on *α*. The somatic boundary condition (11) then results in:

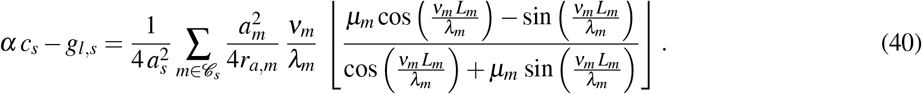

Thus, equations (39) and (40) define a recursive transcendental equation, whose solutions are the constants *α*_*k*_ in (32). NEAT adapts Major’s approach [44] to solve this equation for the *α*_*k*_’s, but replaces the zero finding algorithm with complex analysis techniques based on contour integration [85].

It can be noted that the ratios between all *k*_*n*_ and *k*_*m*_ (*m ∈ C*_*n*_) are fixed by (37), and therefore that they are determined up to a global constant. Major [44] shows that for a delta-pulse input at a location *ξ* ^*′*^, this global constant is *γ*_*k*_*ϕ*_*k*_(*ξ* ^*′*^), with *γ*_*k*_ a factor that is analytically computable once *α*_*k*_ is known. Therefore, it follows that:

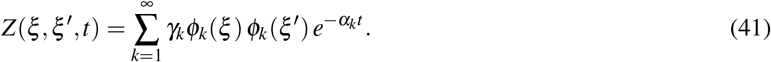

NEAT adopts the convention 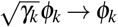 when returning these functions (for instance through `SOVTree.get_important_modes(` thus yielding (32).

### 5.3 The mathematics underlying compartmental neuron models

#### 5.3.1 Neurons as systems of coupled compartments

A compartmental neuron model is governed by a system of equations of the form

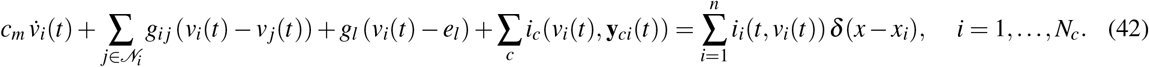

Here, the spatial coordinate 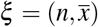 is replaced by a compartment index *i*, and parameters such as capacitance *c*_*m*_, leak *g*_*l*_, or maximal channel conductance are not provided per unit membrane area, as in the morphological models, but simply as a pure capacitance or conductance associated with the compartment. For notational clarity we have omitted the compartment index from these parameters, but in NEAT they are specified on a per-compartment basis. Comparing this equation with the cable model (4), it can be seen that the 2nd-order spatial derivative is replaced by a sum of coupling terms *g*_*i j*_ (*v*_*i*_(*t*) − *v* _*j*_(*t*)), where *g*_*i j*_ is the coupling conductance between compartments *i* and *j*, and where 𝒩_*i*_ denotes the set of neighbour compartments to *i*. Compartmental models in NEAT can be derived in two ways: *(i)* through the 2nd-order finite difference approximation, capturing the full morphology in a detailed manner, and *(ii)* through Wybo’s simplification algorithm, yielding reduced compartmental models.

#### 5.3.2 2nd-order finite difference approximation

Given an equispaced grid {*x*_*i*_} with spacing Δ*x*, the finite difference approximation to the spatial 2nd-order derivative is given by:

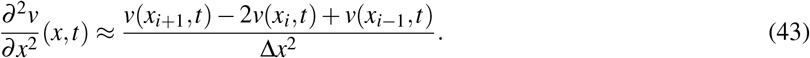

Substituting this in (4) leads to a model of the form (42), where the relation between between cable parameters and compartment parameters has a straightforward interpretation [86]: membrane parameters – defined per unit area – are multiplied with a cylindrical surface, i.e 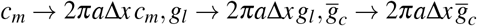, and coupling conductances between compartments are proportional to cylindrical cross section and inversely proportional to inter-compartment distance, i.e. 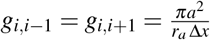(Fig S1A).

At the junction of compartments, NEAT uses a discretization scheme that has empirically been observed to converge to the same parameter values as those yielded by the simplification toolchain (Fig S1B-D). Note that this scheme differs from the schemes implemented by NEURON and/or Arbor. To understand this discretization scheme, it is useful to note that the boundary condition (9), for a single child cylinder 𝒞_*n*_ = {*m*} and equal parameter values in nodes *n* and *m*, approximates the 2nd-order derivative

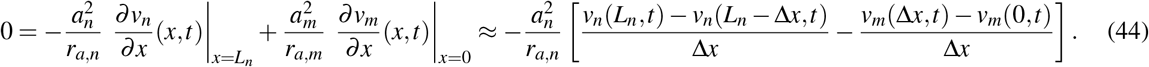

This can be seen from (8), which stipulates that *v*_*n*_(*L*_*n*_, *t*) *≡ v*_*m*_(0, *t*). Note that this equation also models a compartment with zero total current (i.e. no capacitave, leak and channel currents). The geometrical interpretation can then be leveraged, assigning current to this compartment by taking half of the currents on the cylindrical surfaces between the adjacent compartments, yielding e.g. a leak that corresponds to *g*_*l*_ = *πag*_*ln*_ + *πag*_*lm*_. For equal parameters in nodes *n* and *m*, this corresponds to the normal finite difference value 2*πag*_*l,n*_. NEAT’s discretization scheme is therefore as follows: given a maximum inter-compartment distance Δ*x*_max_, NEAT finds the maximal spacing Δ*x* ⩽ Δ*x*_max_, so that the minimum amount of compartments can be distributed on node *n* in an equispaced manner at locations *x* = Δ*x*, 2Δ*x*,…, *L*_*n*_ − Δ*x, L*_*n*_. Coupling conductances between compartments are computed as 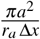, and each compartment receives half the membrane currents that are on the cylindrical surfaces between the compartment and its neighbours (e.g. *g*_*l*_ = *πa*_*n*_ *g*_*ln*_ + ∑_*m∈Cn*_ *πa*_*m*_ *g*_*lm*_ for the leak conductance at a bifurcation).

#### 5.3.3 The simplification toolchain to obtain reduced compartmental models

NEAT implements Wybo’s reduction methodology [31], which is now extended to included concentration mechanism parameters. Given a set of compartment locations {*ξ*_*i*_ | *i* = 1, …, *N*_*c*_} on the morphology, our fit process consists of the following steps: (1) fit the passive leak and coupling conductances, (2) fit the capacitances, (3) fit the maximal conductances of the ion channels, (4) fit the concentration mechanism parameters and (5) fit the leak reversal potentials to obtain the same resting membrane potentials as in the biophysical model.

**Figure S1.**
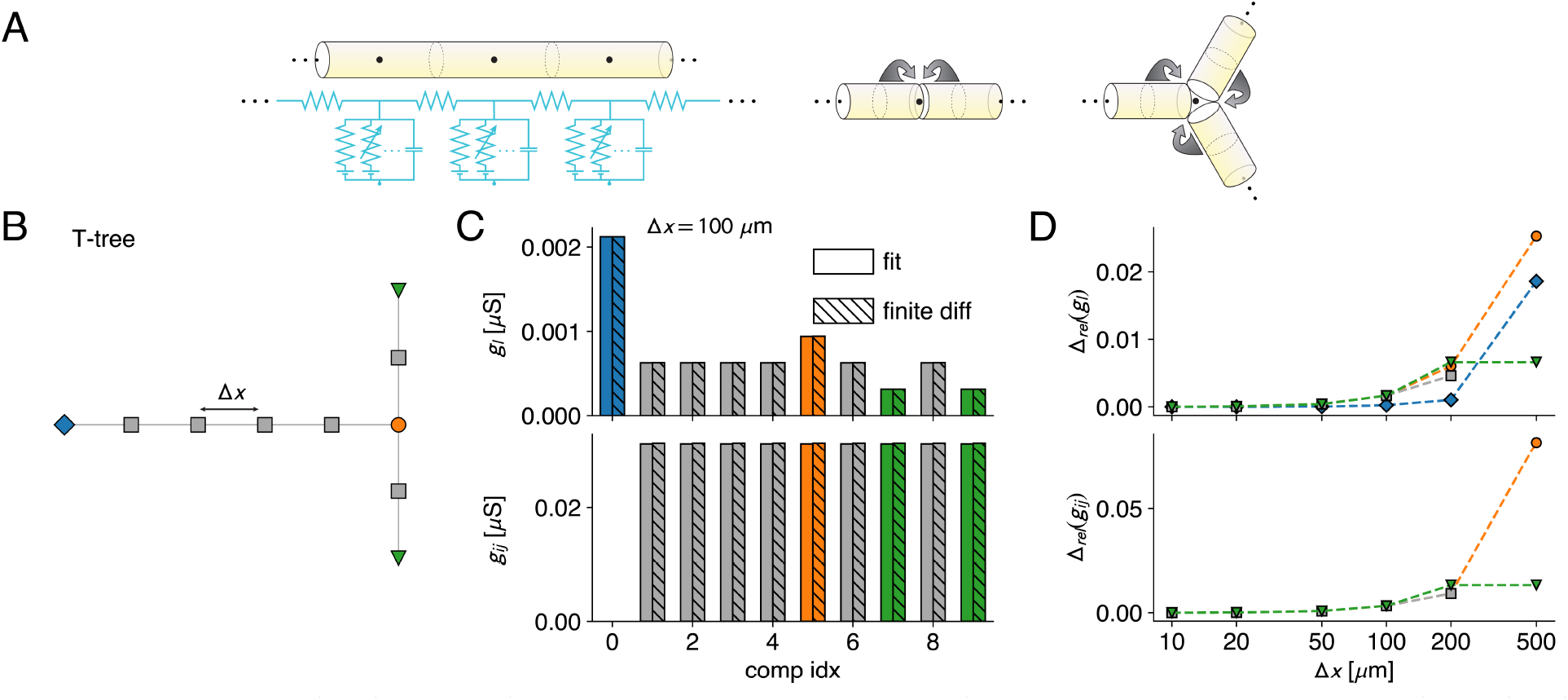
NEAT’s 2nd order finite difference approximation. **A:** In the geometrical interpretation of the 2nd order finite difference approximation, cylinders are subdivided into equally spaced sections, where the effective compartment locations in the are in the middle of each cylinder (left). Compartmental currents are computed as the product of current density and cylinder surface. An interpretation is that each compartment location is assigned half the current density on the cylindrical segments separating it from its neighbours (middle). This scheme is extended for bifurcation compartments (left). Coupling conductances are inversely proportional to the separation between compartments. **B:** Demonstration of the principles of the discretization on a simple T-morphology, featuring a soma (blue diamond) and a single dendrite that branches at a bifurcation (orange circle), terminating in two leaf compartments (green triangles). **C:** NEAT’s discretization scheme converges between compartmental models derived through the reduction methodology (plain bars) and its 2nd order finite difference approximation (hatched bars). The leak conductance (top) at the bifurcation compartment (orange) is 3/2’s the conductance of a normal compartent (grey), and 1/2 that of a normal compartment at the leafs, in agreement with the geometrical interpretation (A). Coupling conductances (bottom) in this scheme are equal for all compartments, as they are equally spaced, and again agree between reduction and finite difference approximation. **D:** Relative difference in parameter values between fit (*p*_fit_) and finite difference (*p*_fd_), i.e. Δ_rel_(*p*) = 2|*p*_fit_ − *p*_fd_|*/*(*p*_fit_ + *p*_fd_), as a function of separation between compartments. Colors indicate compartment type (soma, on a linear dendrite, at bifurcation, or leaf compartment) as in B, C.

1. To fit the passive leak and coupling conductances of the reduced model, NEAT first computes equilibrium voltage and concentrations throughout the detailed model, and creates a passified copy of the model. For this passified model, NEAT computes the matrix 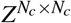, where 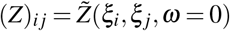. Note that this matrix relates steady-state input perturbation 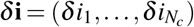 to voltage response 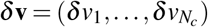, i.e. *δ* **v** = *Zδ* **i**. The fit then determines the admissible coupling terms between compartment sites from the original morphology, and creates a parametric matrix *G* of leak and coupling conductances. This matrix relates voltage response to input current, i.e. *Gδ* **v** = *δ* **i**. Thus, for a perfect fit, *G* should be the inverse of the inverse of *Z*, and therefore *ZG ≈* 𝕀, with 𝕀 the identity matrix, is the fit objective, which NEAT solves in the linear least-squares sense.
2. NEAT fits the capacitances by matching the largest response time-scale of the full model, obtained from (32), to the largest response time-scale of the reduced model.
3. NEAT fits the maximal conductances of each ion channel type separately. To that purpose, NEAT creates copies of the original model where all but one of the channels are removed. For each of these channels *c*, and for a number (*K*) of voltage and state variable expansion points, NEAT then computes the *Z*_*c*1_, …, *Z*_*cK*_ under the quasi-active approximation. If the state variable depends on the Ca^2+^-concentration, NEAT substitutes the equilibrium concentration. NEAT then uses quasi-active approximations in the compartmental model, around the same expansion points as in the full model, to obtain *K* conductance matrices. These matrices can be decomposed as the sum of *G*_pas_, a matrix containing the passive conductances fitted in step 1, and a diagonal matrix *G*_*c,k*_, containing on its diagonal the quasi active terms associated with channel *c*, which in turn feature the to be fitted maximal conductance parameters for each compartment. The fit objective is given by (*G*_pas_ + *G*_*c,k*_*) Z*_*c,k*_ *≈* 𝕀, and is again solved in the linear least-squares sense.
4. Concentration mechanisms, specifically Ca^2+^, are typically described by an equation of the form 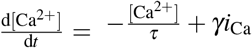, with [Ca^2+^] the Ca^2+^-concentration, *τ* the decay time constant of this concentration, *γ* a fit parameter influencing how strongly Ca^2+^-channels change the Ca^2+^-concentration, and *i*_Ca_ the total current through all Ca^2+^-channels. NEAT fits the *γ*_red_-parameter of the Ca^2+^ in the reduced model to achieve the same concentration in the reduced compartment as at the corresponding site in the mophological model. To that purpose, NEAT sets 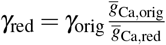, with 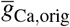 the total maximal conductance densitgy of all Ca^2+^-channels at the compartment site in the full model, and 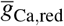 the total maximal conductance of the Ca^2+^-channels at the reduced compartment, which was determined in step 3.
5. Finally, NEAT fits the leak reversals *e*_*li*_ for each compartment *i* = 1, …, *N*_*c*_ to reproduce the equilibrium voltage *v*_*eq*_(*ξ*_*i*_) at the corresponding compartment site in the full model. To achieve this, NEAT evaluates all voltages, concentrations and channel state variables in the full model at equilibrium at all compartment sites, substitutes them in (42) while setting input current *i*_*i*_ and temporal derivative 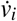 to zero. In this way a system with *N*_*c*_ equations is obtained that is linear in the *N*_*c*_ uknowns *e*_*li*_. This system is then solved using standard Gauss-Jordan elimination.

### 5.4 Simulating the neuron models

Neither NEURON nor Arbor provide direct access to set compartmental parameters; while all simulations of morphological neurons are based on the compartmental description (42), the precise spatial discretization scheme is implemented by the simulator itself. For instance, the NEURON book [1] suggests that maximal compartment spacing should be determined as a user-defined fraction of the frequency dependent length constant *λ* (*ω*), evaluated at 100 Hz. In these simulators, one can either read in morphologies, which are then internally mapped to compartments, or construct neuron models as concatenations of cylindrical sections, which are then discretized internally. Model exports to simulators in NEAT are built to maximize internal consistency with the various analytical frameworks. To that purpose, NEAT proposes two ways of exporting models: either as systems of coupled cylinders – which are most consistent with assumptions in the morphological neuron models, and are then discretized internally by the simulator – or directly as compartmental models – for which the exact compartmental parameters are defined by NEAT.

#### 5.4.1 Exporting morphological models to NEURON

To export morphological models to NEURON, NEAT implements the `NeuronSimTree` class as a subclass of `PhysTree`. Based on the geometrical and physiological parameters stored at each node, a cylindrical `Section` is created in NEURON, that is then connected to its parent section. This cylindrical section is by default discretized following NEURON’s heuristic based on the frequency-dependent length constant *λ* (*ω* = 100 Hz).

#### 5.4.2 Exporting compartmental models to NEST and NEURON

NEAT implements functionality to export compartmental models to both NEST and NEURON. NEST was recently extended with a compartmental modelling framework [69], where custom ion channels and synaptic receptors can be defined through the NESTML model description language [68, 71]. As opposed to NEURON, this compartmental modelling framework exposes the parameters of single compartments. Therefore, the `NestCompartmentTree` is a straightforward extension of the `CompartmentTree`, which instantiates the compartmental model in NEST. The NEURON implementation, however, is somewhat more involved. It requires the creation of a cylindrical section for each compartment, whose surface area, length, membrane parameters, and axial resistance are chosen such that, when this cylindrical section is forced to contain only a single compartment, the compartmental parameters computed internally in NEURON will correspond with those of NEAT’s `CompartmentTree`. These operations are implemented in the `NeuronCompartmentTree`. Note that NEAT has two ways of constructing compartmental models: through the simplification procedure implemented by the `fit_model()` function of `CompartmentFitter`, or through NEAT’s 2nd order finite difference scheme, implemented by the `create_finite_difference_tree()` function of `PhysTree`. therefore, even though NEST’s compartmental modelling framework has a low-level user interface, it can still be used to simulate highly detailed morphological models.

## Notes

### Competing Interest Statement

The authors have declared no competing interest.

